# Normative Sex Differences in Cognition and Morphometric Brain Connectivity: Evidence from 30,000+ UK Biobank Participants

**DOI:** 10.1101/2022.10.12.511938

**Authors:** Crystal C. Yang, Jana F. Totzek, Martin Lepage, Katie M. Lavigne

**Author notes:** Corresponding author. F1128.1, 6875 boulevard Lasalle, Frank B. Common Pavilion, Douglas Mental Health University Institute, Montreal, QC, H4H 1R3, Canada.

## Abstract

There is robust evidence for sex differences in domain-specific cognitive performance in the general population, where females typically show an advantage for verbal memory (VM), while males tend to perform better on tasks of spatial memory (SM). Sex differences in brain structure and connectivity are also well-documented and may provide insight into sex differences in cognition. In this study, we examined sex differences in cognition and morphometric brain connectivity of a large healthy sample (N = 31,180) from the UK Biobank dataset. Using T1-weighted magnetic resonance imaging (MRI) scans and regional cortical thickness values, we applied jackknife bias estimation and graph theory to obtain subject-specific measures of morphometric brain connectivity, hypothesizing that sex-related differences in brain network global efficiency, or overall connectivity, would underlie observed cognitive differences. As predicted, females demonstrated better VM performance and males showed an advantage in SM. Females also demonstrated faster processing speed, with no observed sex difference in executive functioning. Males tended to have higher global efficiency, as well as higher regional connectivity (nodal strengths) in both the left and right hemispheres relative to females.

Furthermore, higher global efficiency in males was found to mediate observed sex differences in cognition, predicting poorer verbal memory performance, better spatial memory, and slower processing speed in males. These findings contribute to an improved understanding of the way biological sex and differences in cognitive performance are related to morphometric brain connectivity as derived from graph-theoretic methods.

---

There is robust evidence for sex differences in cognitive performance. Females perform consistently better in verbal memory tasks (e.g. Bleecker et al. 1988; Kramer et al. 1988; Weiss et al. 2003; Hamilton 2008; Sundermann et al. 2016), whereas males tend to perform better on spatial memory tasks (e.g. Weiss et al. 2003; Postma et al. 2004; Piber et al. 2018). Females therefore tend to show a generalized verbal advantage in their cognitive performance, whereas males show a spatial advantage. In contrast, there is little support for sex differences in executive functioning (Grissom and Reyes 2019), a broad set of cognitive processes which allow an individual to engage in goal-oriented behaviours (Yuan and Raz 2014).

Cognitive functioning is driven by coordinated activity taking place across interconnected and complex brain networks (Bassett et al. 2009; Petersen and Sporns 2015) and individual variations in brain network organization have been found to predict differences in cognitive performance (van den Heuvel and Fornito 2014). Therefore, analyzing brain networks can provide insight into the abnormal organizational dynamics of the brain which may underlie the neurocognitive differences associated with sex (Gong et al. 2009). Examining the connectivity of the brain is likely more informative than univariate analyses of brain regions in developing an understanding of the neurobiology which underlies cognition and behaviour (Seitz et al. 2021; Khalil et al. 2022). Brain connectivity is essential for cognition, such that disruption of these neuronal communications indexed by connectivity leads to signs of cognitive dysfunction (Sporns et al. 2005; Petersen and Sporns 2015). Studying differences in brain connectivity could in this sense give us a fuller picture of brain variations underlying differences in cognitive performance, and one which is likely more informative than studying regional differences alone.

## Brain Connectivity and Structural Covariance

There are many approaches to examining brain connectivity, but especially relevant to understanding sex differences in brain network characteristics are *graph-theoretical* methods. The mathematical approach of graph theory is one of the most common ways of measuring brain connectivity, where the brain is modeled as a complex network consisting of nodes and edges representing the structural or functional connections between regions of the brain (Bullmore and Sporns 2009; Gong et al. 2011; Petersen and Sporns 2015). *Nodes* often represent macroscopic cortical regions, and *edges*, which describe the connections between nodes, may represent white matter architecture (structural connectivity), correlations between regional activity over time (functional connectivity) or correlations between regional structure (e.g., cortical thickness) across individuals (morphometric connectivity) (Sporns et al. 2005; Bullmore and Sporns 2009; He and Evans 2010). The graph measure of nodal *strength* describes the sum of edges that link a particular node to other nodes in a weighted network (Al-Shargie et al. 2019), allowing for an evaluation of the information flow within the network. *Global efficiency* characterizes shorter paths between nodes (Bullmore and Sporns 2009; Alexander-Bloch, Vértes et al. 2013), with *paths* being defined as the minimum number of edges needed to travel between any pair of nodes (Bullmore and Sporns 2009). In this sense, global efficiency represents a measure of how efficiently information is communicated within a network (He and Evans 2010).

*Structural covariance* is a measure of anatomical or morphometric brain connectivity in that it describes correlations between subjects on structural measures, such as volume, cortical thickness (Evans 2013). Inter-individual differences in the structure of brain regions often covary with inter-individual differences in other regions, such that the cortical thickness of one area may influence the thickness of structurally- and functionally-connected brain regions (Alexander-Bloch, Giedd et al. 2013; Alexander-Bloch, Raznahan et al. 2013). For example, many brain regions known to be directly connected by white matter tracts show strong morphological covariation (Alexander-Bloch, Giedd et al. 2013). These correlated brain areas often form networks that have been observed to subserve specific behavioural or cognitive functions (Alexander-Bloch, Giedd et al. 2013). Structural covariance thus represents region-to-region correlations across individuals which could indicate either direct white-matter connections, such as those derived from diffusion-based connectivity methods, or indirect connections through other brain regions, akin to the synchronous activation which is measured in functional connectivity (Bullmore et al. 1997; Rubinov and Sporns 2010; Evans 2013). Although the biological meaning of structural covariance remains debated, it has been proposed to reflect the neurodevelopmental coordination of brain areas (Alexander-Bloch, Raznahan et al. 2013). Measures of structural covariance are calculated by correlating morphological measures, such as cortical thickness, surface area, or regional volumes of brain regions, across a group of subjects and examining differences between groups (Evans 2013; Carmon et al. 2020). Structural covariance of cortical thickness, which reflect the density and arrangement of cells, including neurons, neuroglia, and nerve fibers (Parent and Carpenter 1996; Narr et al. 2005) may be of particular interest as it has been linked to underlying structural connectivity (Chen et al. 2008; Gong et al. 2012).

## Normative Sex Differences in Brain Connectivity

In addition to differences in cognition, there is also evidence for sex differences in structural brain connectivity (e.g. Lv et al. 2010; Gong et al. 2011). In a graph-theoretic analysis of *diffusion tensor imaging* (DTI) tractography, Gong and colleagues (Gong et al. 2009) found female sex to be related to greater overall cortical connectivity, as well as higher local and global efficiency in cortical network organization, suggesting a more efficient use of available white matter. The authors furthermore observed hemispherically-asymmetrical sex differences related to regional network efficiency, such that females had higher efficiency in regions within the left verbal-dominant hemisphere, and males had higher efficiency in regions of the spatial-dominant right hemisphere. Furthermore, using DTI methods, Ingalhalikar and colleagues (Ingalhalikar et al. 2014) found that whereas male brains showed greater intrahemispheric connectivity, female brains rather tend to show higher interhemispheric connectivity. Female sex has also been found to predict stronger coupling between structural-functional connectivity patterns (Zhao et al. 2021). Additionally, in a study examining functional network activity, Tian and colleagues (Tian et al. 2011) found that males had greater *clustering*, where specific groups of nodes are highly connected to each other (Bullmore and Sporns 2009), in the right hemisphere, but lower clustering in the left hemisphere, suggesting asymmetrical sex differences in functional network topological organization. Despite these studies, little work has examined sex differences in morphometric brain connectivity. Though differences in structural covariance have been related to biological sex (e.g. Gong et al. 2009; Abbs et al. 2011; Persson et al. 2014), such differences, which are based on sex-aggregated data, cannot be subsequently related to subject-specific measures, such as cognitive performance. In this study, we addressed this limitation by applying the Jackknife bias estimation procedure (Das et al. 2018; Ajnakina et al. 2021) to compute subject-specific morphometric brain connectivity measures and allow for the investigation of brain connectivity underlying sex differences in cognition.

## Specific Aims and Hypotheses

The goal of this study was to examine normative sex differences in cognition and brain connectivity. We examined a sample derived from the UK Biobank data repository (UKBB; https://www.ukbiobank.ac.uk/) and examined sex differences in cognition and cortical thickness-derived morphometric connectivity. Then, by employing subject-specific morphometric connectivity, we were able to assess the extent to which brain connectivity mediated sex effects on cognition. We first tested the hypothesis that there are normative sex differences in domain-specific cognitive performance, such that females would show a verbal memory advantage and males would exhibit a spatial memory advantage, with no expected sex differences in executive functioning. Due to the mixed nature of previous findings, we took an exploratory approach to examining sex differences in global and local brain connectivity. Further, we predicted that differences in global brain connectivity would mediate observed differences in cognitive performance.

## Methods and Materials

### Participants

Participants were acquired from the UKBB dataset (https://www.ukbiobank.ac.uk/), a large-scale database containing a diverse range of health data belonging to over 500,000 adult individuals living in the United Kingdom (Sudlow et al. 2015). The set of data available in the UKBB includes genetic, MRI, and sociodemographic information. The dataset is open access to the global research community, and research applications must be approved by an independent access subcommittee (Sudlow et al. 2015). Participants completed behavioural testing, MRI scans, and the collection of biological samples (Sudlow et al. 2015). For this study, we worked with data from 31,080 participants with T1 structural MRI scans collected at a follow-up assessment.

General exclusion criteria accounting for MRI safety and quality standards were applied. More precisely, subjects were included if they weighed less than the scanner table load limit of 170 kg and did not have contraindications including electrical or metal implants (eg. a cardiac pacemaker or ferromagnetic implants), recent surgery in the last six weeks, hearing difficulties, breathing difficulties, tremors, claustrophobia, or pregnancy (Miller et al. 2016; Machann et al. 2020; UKBB). Participants were also included if they had no history of substance abuse, had never been diagnosed with a mental and behavioural disorder (ICD-10 Chapter V; F00 - F99), did not have any neurological conditions (ICD-10 Chapter VI; G00 - G99), and had never experienced any instances of traumatic brain injury (ICD-10; F06, F07). Group differences in the Index of Multiple Deprivation (IMD), a holistic measure of socioeconomic deprivation, were also examined. This pre-analysis assessment accounted for the fact that socioeconomic status and education are hypothesized to contribute to *cognitive reserve*, or the increased capacity and functioning of existing brain networks which is protective against age- or disease-based brain pathology and corresponding cognitive performance decline (Barulli and Stern 2013). We included only participants with Index of Multiple Deprivation (IMD) England scores, as this comprised the vast majority of participants and removed the need to scale regionally-normalized IMD data from Wales and Scotland.

### Behavioural Methods

#### Neuropsychological Assessments (UKBB Category 100026)

##### Verbal Memory

A ‘paired associate learning task’ was used to assess verbal memory (https://biobank.ndph.ox.ac.uk/ukb/label.cgi?id=506). Participants were shown 12 pairs of words for a total of 30 seconds. After an interval in which they completed a separate unrelated test, participants were then presented with the first word of ten of these pairs and asked to select the matching second word from four given choices. The number of word pairs correctly associated is used to calculate this variable, with scores ranging from zero to a maximum of ten correctly associated pairs. Fawns-Ritchie and Deary (Fawns-Ritchie and Deary 2020) found this task to have good validity (*r* = 0.46) and moderate reliability (*r* = 0.45).

##### Spatial Memory

Participants completed a visuo-spatial memory task, labeled ‘pairs-matching’ (https://biobank.ndph.ox.ac.uk/ukb/label.cgi?id=100030). In this task, participants were asked to memorize the position of matching pairs of cards on a screen. The cards were then turned face down, and participants were asked to touch as many matching pairs as possible in the fewest number of tries. Two rounds of this task were conducted, with the first round featuring three pairs of cards and the second featuring six pairs. The number of incorrect matches in a round is used to represent this variable. This task has been found to show moderate concurrent validity (*r* = −0.29) and reliability (*r* = 0.41) (Fawns-Ritchie and Deary 2020).

##### Processing Speed and Executive Function

Processing speed and executive function were assessed using a ‘trail making’ task (https://biobank.ndph.ox.ac.uk/ukb/label.cgi?id=505). In this task, participants were presented with a series of digits (numeric trial) or digits and letters (alphanumeric trial) located within circles scattered around the screen. The *numeric trail-making task* consists of only numbers which should be selected in an ascending pattern, and is thought to reflect baseline processing speed, in the form of motor and visual search speed abilities (Arbuthnott and Frank 2000; Delis et al. 2012). In contrast, the *alphanumeric trail* consists of both letters and numbers which must be selected in an alternating pattern according to their order, and is believed to measure more general executive functioning, specifically set-shifting and cognitive flexibility (Arbuthnott and Frank 2000; Kortte et al. 2002). The duration of time taken to complete both paths was used to assess performance on this task, with performance on the numeric trail reflecting processing speed and performance on the alphanumeric trail reflecting executive function. Whereas numeric trail-making is hypothesized to directly measure processing speed (Laere et al. 2018), some work has suggested that alphanumeric trail-making performance, and higher-order executive functions in general, are also likely highly dependent on processing speed (Salthouse 2011; MacPherson et al. 2017). The numeric task demonstrates moderate validity (*r* = −0.31) and good reliability (*r* = 0.48), whereas the alphanumeric task demonstrates good validity (*r* = −0.50) and reliability (*r* = 0.77) (Fawns-Ritchie and Deary 2020).

### Neuroimaging Methods

#### MRI Acquisition Type

All MRI scans in the dataset were performed on a single scanner with a Siemens Skyra 3T scanner and a Siemens 32-channel RF receive head coil (Smith et al. 2020). For our analysis, we used high-resolution T1-weighted structural MRI scans (resolution = 1 x 1 x 1 mm voxels, field-of-view = 208 x 256 x 256 matrix, acquisition time = 5 min, in-plane acceleration factor (R) = 2, TI/TR = 4880/ 2000 ms) (Alfaro-Almagro et al. 2018; Smith et al. 2020). Scans were taken with a three-dimensional magnetization-prepared rapid acquisition with gradient-echo (3D MPRAGE), Integrated Parallel Acquisition Technique (iPAT) = 2, and the “prescan-normalize” signal normalization filter (Smith et al. 2020).

#### Image Processing

Scans were preprocessed through the Corticometric Iterative Vertex-based Estimation of Thickness (CIVET) pipeline (Ad-Dab’bagh et al. 2006) through the NeuroHub platform (https://neurohub.ca) using CBRAIN (Sherif et al. 2014). The automated CIVET pipeline estimated cortical thickness at 40962 points for each hemispheric surface throughout the cortex (Collins et al. 1994; Rajaprakash et al. 2014). Each image was standardized through linear registration to the MNI ICBM Average Brain Stereotaxic Registration Model (Collins et al. 1994). A non-uniformity correction algorithm was applied to the images (Sled et al. 1998), and brain tissue was classified into cerebrospinal fluid, grey matter, and white matter, with the background differentiated from the tissue (Zijdenbos et al. 2002). The Constrained Laplacian Anatomic Segmentation using Proximity (CLASP) algorithm was used to extract cortical surfaces (Kim et al. 2005). Non-cerebellar brain structures, including the brain stem and the cerebellum, were removed (Lv et al. 2010). The Euclidean distance between corresponding nodes of the white and grey matter surfaces was measured to calculate the cortical thickness estimates (Lerch and Evans 2005). Participant scans were processed through an automated machine-learning quality control procedure developed in-house (https://github.com/joshunrau/civetqc/tree/main/civetqc/; Unrau et al. 2022) to exclude problematic scans. After removing scans that did not meet criteria, our final sample size was 31,080 participants. Lastly, the surface-based Desikan-Killiany Tourville (DKT) cortical labeling protocol was used to parcellate the regions of interest and obtain mean regional cortical thickness values for each individual (Klein and Tourville 2012).

#### Jackknife Bias Estimation Procedure

We first established the full-sample structural covariance (Alexander-Bloch, Giedd et al. 2013; Alexander-Bloch, Raznahan et al. 2013; Evans 2013) between DKT regions using pairwise correlations. Subject-specific structural covariance matrices were then calculated using the jackknife bias estimation procedure (https://github.com/katielavigne/jackknife_connectivity). This bootstrapping method employs a “leave-one-out” approach by recalculating the structural covariance matrix N times, for each iteration of N-1 participants (Miller 1974; Richter et al. 2015; Das et al. 2018; Ajnakina et al. 2021). Subtraction of these leave-one-out structural covariance matrices from the full-group covariance matrix then allows for the determination of a single subject’s contribution to the overall group-level covariance structure (Das et al. 2018, S3-S4).

#### Morphometric Brain Connectivity

We used weighted graphs, which may be preferable to binary graphs as a representation of brain connectivity, since binary graphs are based on thresholds that may be arbitrary and inconsistent across groups. As a result, binary graphs may artificially oversimplify connectivity patterns and remove the nuances of individual topological information (Fornito et al. 2013; Yeh et al. 2021). A weighted network representation of the structural connectivity of the brain may therefore be a more biologically informative representation of the properties of this network (Fornito et al. 2013; Bassett and Bullmore 2017; Yeh et al. 2021).

The subject-specific graph measure of *global efficiency* was then computed for each participant through the Brain Connectivity Toolbox in Python (bctpy; Rubinov and Sporns 2010) as a measure of overall connectivity. *Global efficiency* is defined as the average inverse shortest path length of the network and was calculated using the following equation:

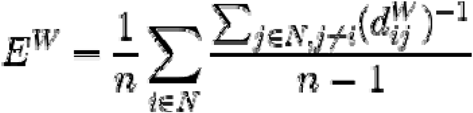

In this equation, *N* represents the complete set of nodes in the graph, *n* indicates the number of nodes, *i* and *j* are any two nodes in the network, and 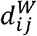 is the shortest path length in the weighted network between nodes *i* and *j* (Rubinov and Sporns 2010). We calculated weighted global efficiency (*E^W^*), which is the measure of global efficiency as calculated within a *weighted graph*. Nodal *strength* (*k_i_^w^* also known as weighted degree, is the sum of weighted edges which are connected to a given node, reflecting nodal, or regional, connectivity in a weighted graph (Bullmore and Sporns 2009). Each node was defined as a single region of the DKT atlas, presented in Supplementary Figure 1. We calculated strength using the following equation:

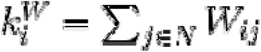

### Statistical Analyses

#### General Linear Models for Cognitive Measures and Global Efficiency

We conducted multiple linear regressions, dummy-coding for sex (female = 0, male = 1). The following general linear model (GLM) equation with the predictor of sex and a covariate of age was applied for each dependent variable of global connectivity, verbal memory, spatial memory, and executive functioning:

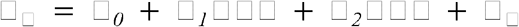

Due to missing data, some participants were excluded from the statistical analyses of our cognitive variables. As a result, our final healthy control sample for these analyses was composed of 27,307 participants for spatial memory and 20,642 participants for all other cognitive variables.

#### Regional Connectivity Analysis

To assess sex differences in regional connectivity, we conducted linear regressions for every nodal strength, using the following linear model.

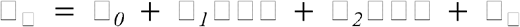

We controlled for Type I error from multiple comparisons using the Bonferroni

Correction method, using an adjusted *p* value (□ / n), where n is the number of comparisons conducted. Since we were comparing the nodal strength of every DKT region in addition to global efficiency and four cognitive variables, giving us sixty-seven comparisons, our corrected *p* value = .0007.

#### Exploratory Mediation Analysis

Significant direct effects involving global efficiency were followed up with a mediation analysis to assess if overall brain connectivity mediates the relationship between sex and cognitive performance. To conduct this analysis, we used the processR package (processR; Moon 2021), which adapted the Hayes PROCESS Macro analysis and modeling tool (Hayes 2018) for usage in R. A schematic summarizing the full methods used in this study is presented as Figure 1. To visualize the graph-theoretical measures of strength and efficiency, BrainNet Viewer (Xia et al. 2013, https://www.nitrc.org/projects/bnv/), was used. The hypothesized mediation model is visually represented by Supplementary Figure 2.

**Figure 1.**
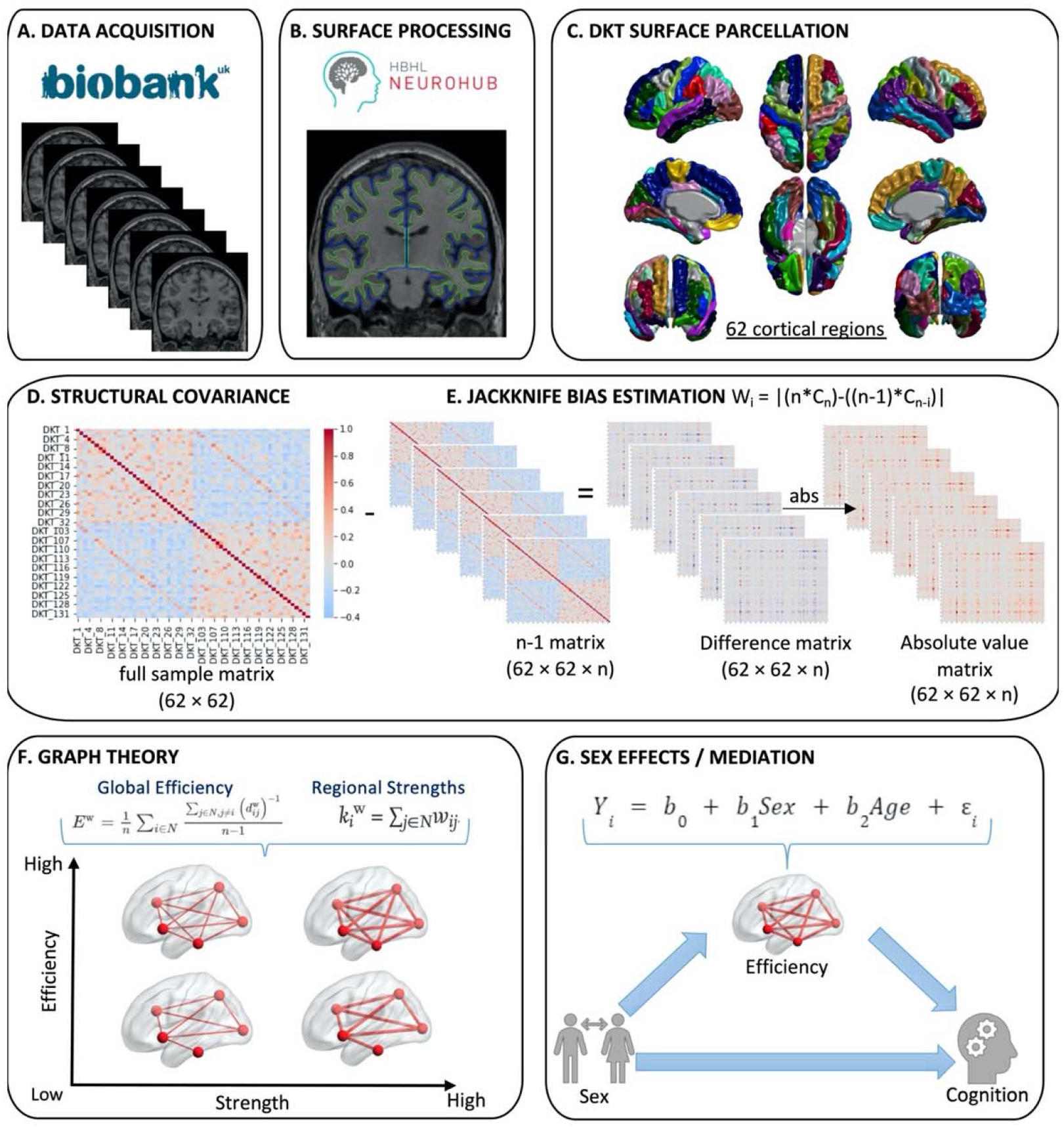
Data processing and analysis schematic. *Note*. A: T1-weighted MRI acquisition, B: NeuroHub processing, C: DKT Parcellation, D: Structural covariance calculation, E. Jackknife Bias Estimation Procedure, F: Graph theory, G: Sex differences analysis. For F. and G. the BrainNet Viewer (Xia et al., 2013, http://www.nitrc.org/projects/bnv/) was used to visualize strengths and efficiencies.

### Data and Code Availability

The analysis code used in this paper is available in the GitHub links specified in the Methods section of this paper. Although the UK Biobank dataset is open access to the global research community, research applications and access to data must be approved by an independent access subcommittee (Sudlow et al. 2015). Access to data for this study was approved under a larger application (ID: 45551) as part of the Healthy Brains for Healthy Lives (HBHL; https://www.mcgill.ca/hbhl/) initiative at McGill University. Data was downloaded through NeuroHub, and access was supported by Calcul Québec and Compute Canada.

### Ethics

The UK Biobank originally received an ethical review and approval from the North West Multi-centre Research Ethics Committee (MREC) in 2011, and approval was renewed in 2021. UK Biobank participants gave informed consent for UK Biobank to collect, store and make available a range of data, as well as to follow their health over time in the interest of health-related research. Ethical approval for this study was under a larger application (ID: 45551) as part of the Healthy Brains for Healthy Lives (HBHL; https://www.mcgill.ca/hbhl/) initiative at McGill University.

## Results

We first conducted simple linear regressions to assess group differences in age, since age is related to performance on all cognitive domains of interest (Moffat et al. 2001; Kramer et al. 2003; Tombaugh 2004) as well as brain connectivity (Fjell et al. 2016). As a significant difference in age was found between groups (*F*(1,31078) = 198.10,*p* < .001, *R*^2^_adj_ = .006), it was included as a covariate. No significant differences in IMD scores were found in our sample, and were therefore not controlled for. Full demographics are summarized in Table 1.

**Table 1.** Sample Characteristics.

| Sample (N = 31080) |  |  |
| --- | --- | --- |
|  | Females<br>(*n* = 16223) | Males<br>(*n* = 14857) |
| % Right-handed | 90.3 | 87.6 |
| Mean Age (years) | 62.8 | 63.9* |
| SD | 7.3 | 7.6 |
| Mean Edu Score | 12.1 | 12.3 |
| SD | 13.3 | 13.7 |
| Mean IMD | 15.3 | 15.2 |
| SD | 11.8 | 12.00 |
| Mean VM Correct | 7.4 | 6.6* |
| SD | 2.5 | 2.6 |
| Mean SM Incorrect | 0.4 | 0.3* |
| SD | 0.8 | 0.8 |
| Mean PS Numeric Time | 216.0 | 225.8* |
| SD | 84.0 | 82.9 |
| Mean EF Alphanumeric Time | 216.0 | 550.4 |
| SD | 257.7 | 261.1 |
| Mean Global Efficiency | 0.8 | 0.9* |
| SD | 0.3 | 0.5 |

*Note.* SD = standard deviation, Edu Score = education score on the IMD England, IMD = Index of Multiple
Deprivation England score, VM Correct = number of correct matches on the verbal memory task, SM Incorrect = number of incorrect matches on the spatial memory task, EF = executive functioning task, PS Numeric Time = time taken to complete numeric path in processing speed task, EF Alphanumeric Time = time taken to complete alphanumeric path in executive functioning task. We controlled for Type I error from multiple comparisons using the Bonferroni Correction method, using an adjusted *p* value ( $\alpha / n$ ), where *n* is the number of comparisons conducted. Since we were comparing the nodal strength of every DKT region as well as four cognitive variables, giving us sixty-seven comparisons, our Bonferroni-corrected *p* value = .0007
\* $p < .0001$

### Sex Differences in Cognition

Testing our hypotheses for normative sex differences, we observed a significant effect of sex on verbal memory (*F*(2, 20639) = 694.30,*p* < .001, *R^2^_adj_* = .06), where females showed a small to medium advantage indicated by more correct word pairs (*B* = 0.80, *t*(20639) = 3.28, *p* < .001). Moreover, a significant effect was found for spatial memory (*F*(2,29304) = 167.30,*p* < .001 *R^2^_adj_* = .01), where males in our sample showed a small but significant advantage, making fewer incorrect matches (*B* = −0.06, *t*(29304) = −6.35, *p* < .001). An effect was also found for processing speed (*F*(2, 20639) = 899.90,*p* < .001, *R^2^_adj_* = .08), where females took less time to complete the numeric trail-making task, indicating an advantage (*B* = −5.95, *t*(20639) = −5.28,*p* < .001). No significant sex differences were found for performance on the alphanumeric trailmaking task. Full results are presented in Table 2 and visually modeled in Figure 2.

**Table 2.** Sex Differences in Sample.

| | *B* | *SE* | *t* | $R^2_{adj}$ |
| --- | --- | --- | --- | --- |
| Sex (male) |  |  |  |  |
| Verbal Memory* | -0.80 | 0.04 | -23.64 | .06 |
| Spatial Memory* | -0.06 | 0.01 | -6.35 | .01 |
| Processing Speed* | 5.95 | 1.13 | 5.28 | .08 |
| Executive Functioning | 5.87 | 3.49 | 1.68 | .07 |
| Global Efficiency* | 0.09 | 0.01 | 20.50 | .03 |

*Note.* *B* = unstandardized beta coefficient, *SE* = standard error, $R^2_{adj}$ = adjusted R squared, VM = number of correct matches on the verbal memory task, SM = number of incorrect matches on the spatial memory task, PS = time taken to complete numeric path in processing speed task, EF = time taken to complete alphanumeric path in executive functioning task, GE = Global Efficiency.
\* $p < .0001$

**Figure 2.**
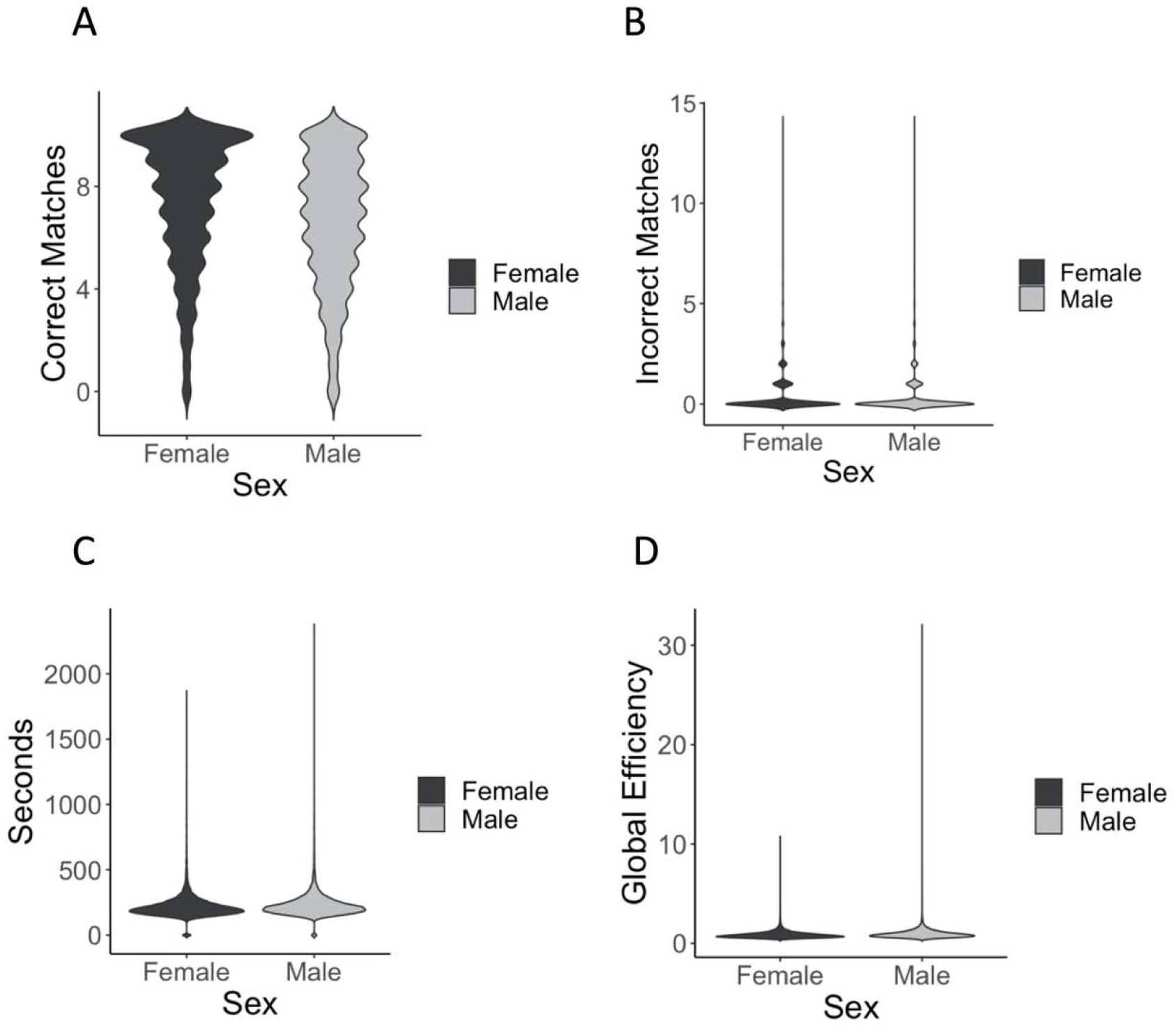
Sex differences on verbal memory (A), spatial memory (B), processing speed (C), and global efficiency (D) *Note*. A: Correct matches on paired associate learning task, B: Incorrect matches on pairs-matching task, C: Seconds to complete numeric path, D: Global efficiency.

### Sex Difference in Connectivity Measures

Sex showed a small but significant effect on global efficiency (*F*(2, 31077) = 526.70, *p* < .001, *R^2^_adj_* = .03), such that it was higher in males (*B* = 0.09, *t*(31077) = 20.50,*p* < .001). As regionally defined by the DKT atlas, we additionally found that males showed higher nodal strengths throughout the brain, with the greatest differences observed in parietal regions and nonsignificant effects in bilateral insula (Figure 3, Supplemental Table 1, *p* < .05, corrected).

**Figure 3.**
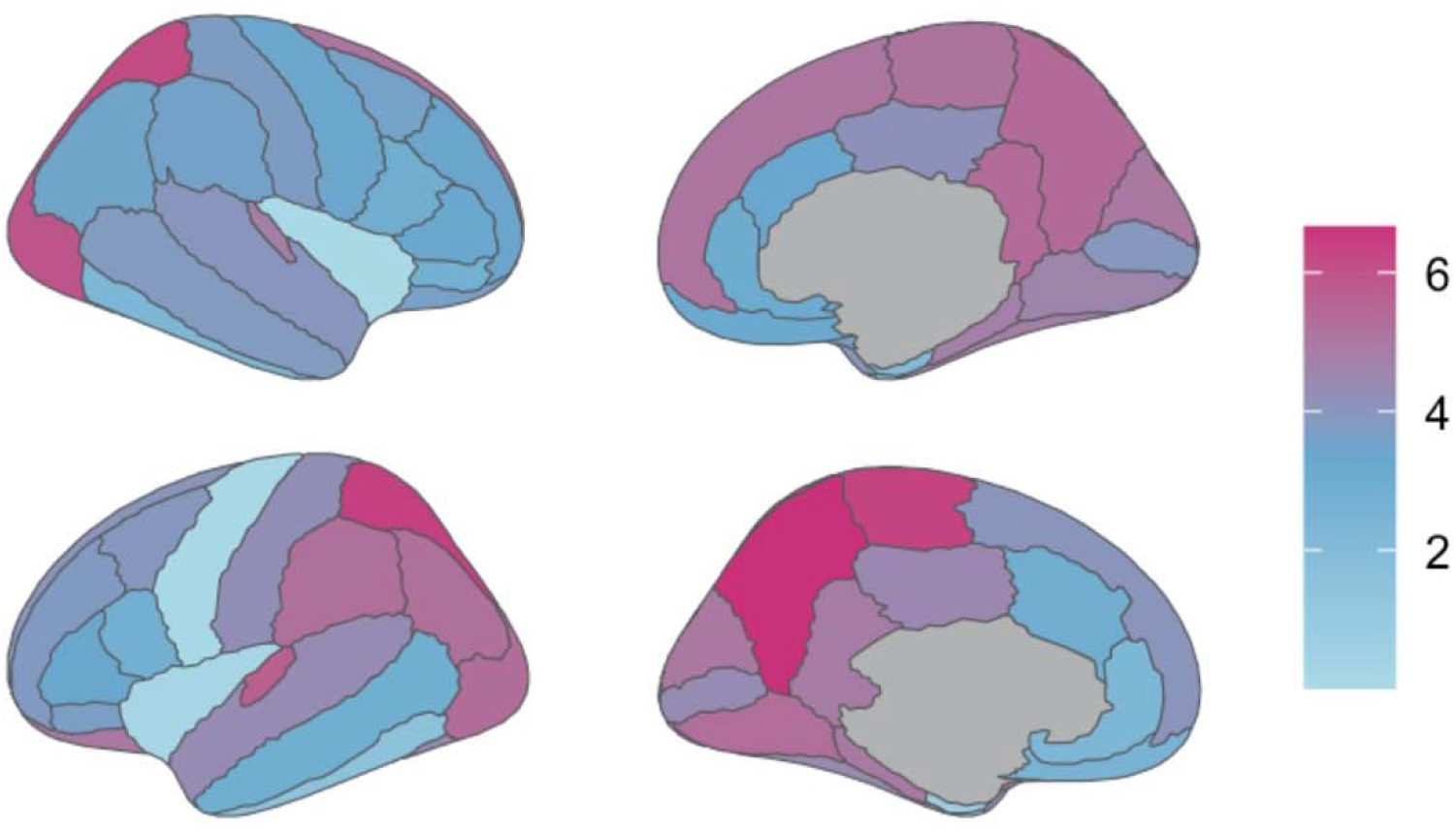
Regional sex differences in morphometric connectivity strength using the DKT atlas. Note. Effect sizes for sex differences in nodal strength by DKT region.

### Mediation Analyses

As presented in Table 3, we ran multiple mediation models to test if global efficiency significantly mediated sex differences in cognition. We found that global efficiency significantly mediated the sex differences in verbal memory (*R^2^* = .06, *F*(3, 20638) = 442.69,*p* < .001, 95% CI [−0.04, −0.02]), spatial memory (*R^2^* = .01, *F*(3, 29303) = 112.85,*p* < .001, 95% CI [0.00, 0.00]), and processing speed (*R^2^* = .08, *F*(3, 20638) = 602.04,*p* < .001, 95% CI [0.06, 0.60]). These results suggest that higher overall connectivity in males contributed to more errors on the verbal memory task, more correct matches on the spatial memory task, and longer processing speed times. The results of these mediation analyses are visually modelled in Figure 4.

**Table 3.** Index of Moderation of Sex Differences by Global Efficiency.

|  | *B* | Bootstrapped SE | 95% Bootstrapped CI |
| --- | --- | --- | --- |
| Verbal Memory* | -0.03 | 0.00 | [-0.04, -0.02] |
| Spatial Memory* | 0.00 | 0.00 | [0.00, 0.00] |
| PS Numeric* | 0.31 | 0.14 | [0.06, 0.60] |

*Note.* Verbal Memory = number of correct matches on the verbal memory task, Spatial Memory = number of incorrect matches on the spatial memory task, PS Numeric = time taken to complete numeric trail-making processing speed task, EF Alphanumeric = time taken to complete alphanumeric trail-making executive functioning task, *B* = unstandardized beta coefficient, SE = standard error, CI = confidence interval. \* $p < .05$

**Figure 4.**
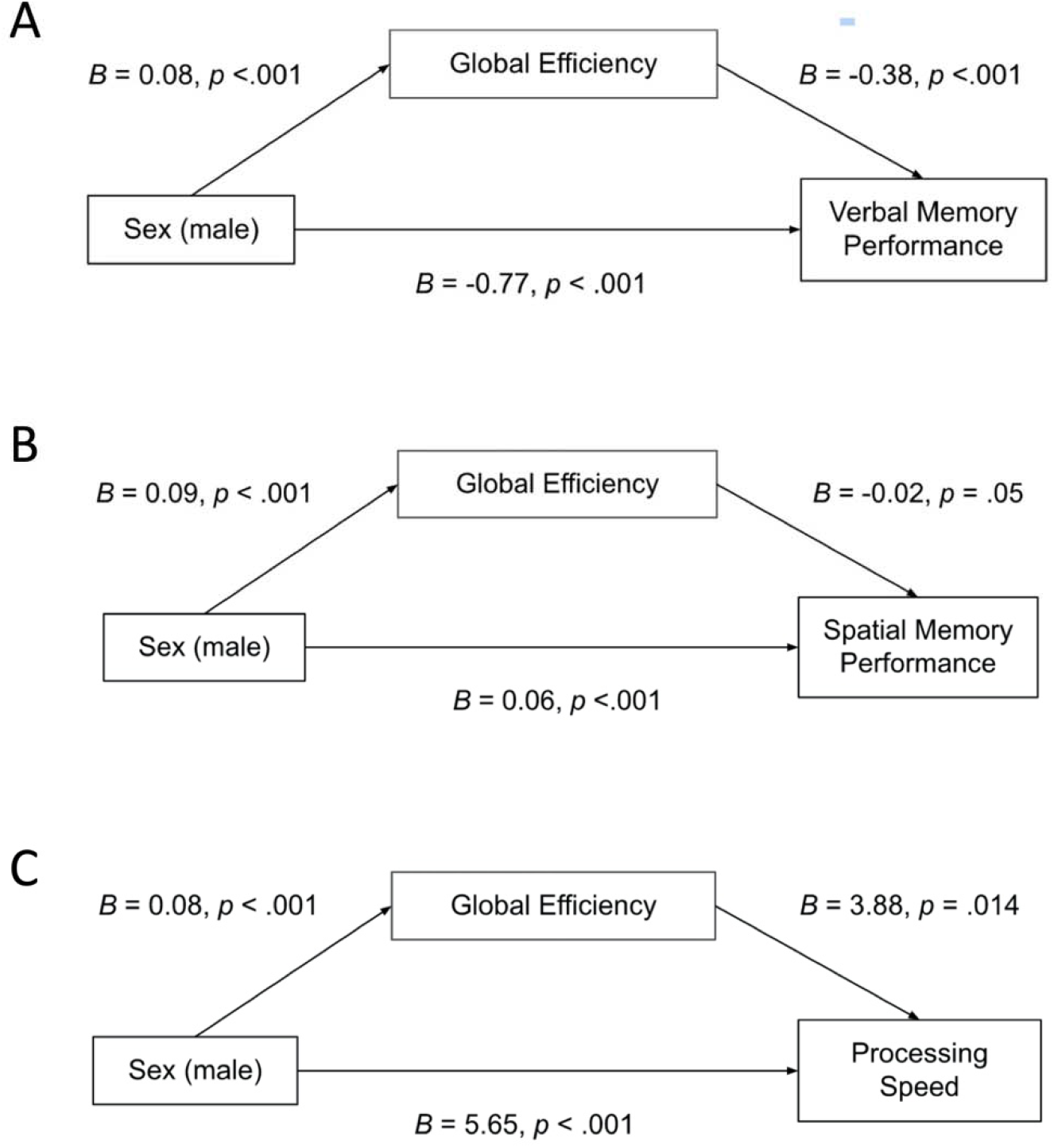
Global efficiency as a mediator of verbal memory (A), spatial memory (B), and processing speed. *Note*. These figures correspond to Model 4 in Hayes PROCESS Macro (Hayes, 2018).

## Discussion

The purpose of this study was to assess well-known sex differences in domain-specific cognitive performance and how these differences might relate to sex differences in individual anatomical brain connectivity. The extraordinary size of our sample (*n* = 31,080) makes it the largest to-date for a study of sex differences in cognitive performance and brain connectivity. Consistent with the literature, females exhibited an advantage for verbal memory while males showed an advantage for spatial memory. Interestingly, females additionally showed an advantage in processing speed, as indicated by performance on the numeric trail-making task. No significant sex difference was observed in executive functioning, as indicated by set-shifting ability measured with the alphanumeric trail-making task. Males in our sample had higher global efficiency than females, as well as higher nodal strengths in both the left and right hemispheres, appearing to demonstrate both greater global and local brain connectivity. Furthermore, global efficiency was found to mediate sex differences in cognition, where higher global efficiency predicted poorer verbal memory performance, better spatial memory, and slower processing speed in males.

The observed sex differences in verbal memory and spatial memory observed in this study are well-documented in previous literature (e.g. Weiss et al. 2003). However, it remains unclear why greater global efficiency in males appears to mediate poorer verbal memory and better spatial memory performance in our sample. One consideration is that the current study on cortical thickness does not account for hippocampal structure or connectivity, a region that is key for memory processes. Well-documented sex differences in hippocampal volume may drive these bidirectional effects. In a functional connectivity study, Spets and colleagues (Spets et al. 2021) found that whereas females did not show greater connectivity than males in any brain regions, males showed greater regional connectivity in the anterior prefrontal cortex, precuneus, cingulate sulcus, and medial posterior frontal cortex, which the same study observed to be functionally connected to the hippocampus during a spatial long-term memory task. Interestingly, in our results, sex effects were less pronounced in the left than right entorhinal cortex, though both still showed greater nodal strength in males. This could suggest lateral differences in the connectivity of this region related to sex. Dysconnectivity between the entorhinal cortex and the hippocampus has previously been related to spatial memory deficits (Parron et al. 2006). Stone et al. (Stone et al. 2011) found that deep brain stimulation promoting neurogenesis of the entorhinal cortex, and subsequent incorporation into hippocampal circuits, facilitated improved spatial memory performance in mice. More studies are needed to understand the significance of sex differences in morphometric connectivity involving the medial temporal lobe and well-established sex differences in memory.

The female advantage in processing speed that was observed in the present study has also been reported in previous work. For example, Camarata and Woodcock (Camarata and Woodcock 2006) noted that females scored higher on measures of processing speed across large standardized samples for three cognitive batteries. Furthermore, in a US-standardized sample (*n* = 2450), Irwing (Irwing 2012) saw that females showed a large processing speed advantage as measured on the WAiS-iii. it remains to be determined why decreased global efficiency appeared to mediate faster processing speed in females in our sample. Imms et al. (Imms et al. 2021) found that better processing speed was correlated with both higher global and network-specific efficiency. These differences in findings signals the need for more studies in order to better understand sex differences in cognition and underlying brain connectivity derived from graph-theoretic methods.

Given the unexpected direction of some results in our study, it is important to note that relatively few studies to date have published results examining sex differences in morphometric brain connectivity as derived from measures of cortical thickness, and existing findings are mixed. For example, Lv et al. (Lv et al. 2010) examined sex differences in structural covariance based on CIVET-derived cortical thickness measures. Although they found two increased interregional correlations related to female sex in their sample (*n* = 184), they concluded that there were no overall differences in local and global connectivity, and that healthy adults did not show significant differences in overall network topology. Studies using diffusion-based connectivity have previously reported either lower levels of global efficiency in males (e.g. Gong et al. 2009) or no sex effects on global efficiency (Yan et al. 2011). While it is difficult to compare findings across modalities, maps created from cortical thickness correlations are strikingly similar to maps resulting from DTI techniques examining white matter tracts, which suggests that cortical thickness correlations are approximately reflective of true neuroanatomical connections (Lerch et al. 2006; Zhang et al. 2012). For example, in a study examining structural covariance in cortical thickness and DTI tractography, Gong and colleagues (Gong et al. 2012) found that approximately 35-40% of thickness correlations converged with tractography findings. These results were taken to suggest that cortical thickness correlations were partially reflective of white matter connections, but also contained unique connectivity information not captured by diffusion methods. Additionally, anatomical networks derived from structural covariance analyses of cortical thickness show small-world topological organizational properties that are consistent with those of networks derived from functional activity (He et al. 2007; Bassett et al. 2008).

Our finding that local connectivity, as quantified by nodal strength, was higher bihemispherically for males also contrasts with previous findings using DTI graph methods. For example, Gong et al. (Gong et al. 2009) observed that females showed increased regional efficiency of areas in the left verbal-dominant hemisphere, being Heschl’s gyrus, the superior temporal gyrus, insula, and superior and inferior parietal gyri. In contrast, males showed higher regional efficiency in only the right rolandic operculum and triangular inferior frontal gyrus. However, these results were obtained from DTI tractography data rather than cortical thickness estimates, and are based on regional efficiency as a measure of local connectivity rather than nodal strength. Similarly, Tian and colleagues (Tian et al. 2011) found a sex difference in asymmetrical hemispheric connectivity such that males had greater right hemispheric connectivity than females, as was predicted in the present study. However, these results were derived from functional MRI data and reflected a measure of clustering rather than nodal strength. Greater investigation is required to understand the meaning of these methodological differences between studies and how they might relate to differences in findings.

### Strengths and Limitations

A notable strength of this study was the healthy sample that we were able to extract from the UKBB which is exceptionally large for a neuroimaging study (Szucs and Ioannidis 2020). We are unaware of any other studies which have conducted graph analyses of morphometric brain connectivity with such a large sample size (N = 31,080). Furthermore, we were rigorous in selecting exclusion criteria, which increases the likeliness that true sex differences were represented in our findings. However, our study was limited by the fact that there was a large amount of missing data for participants, leading to limitations in our analyses, such as the reduction in the sample size of our cognitive analyses. Even so, this was likely mitigated by the large size of our remaining sample. Furthermore, ethnicity information was largely missing from our data, preventing an assessment of group differences.

The significant but small effect size of global efficiency suggests that this finding is reliable in our sample, but may not be of practical relevance. It is possible that the higher global and local brain connectivity found in males in our sample results from an unidentified factor that was not controlled for in our analyses, either giving rise to skewed sample characteristics or influencing our calculations of individual-specific global connectivity. The sample also showed large variation in global efficiency which could suggest an unidentified factor such as participants who were not properly screened by our exclusion criteria for mental or behavioural disorders, substance abuse disorders, neurological conditions, neurodegenerative diseases, or traumatic brain injuries, due to these conditions being unreported or undiagnosed. Our findings may also be influenced by the fact that participants in this UKBB sample were generally older adults (*mean age* = 63.5 years). Results from this sample pertaining to global efficiency may therefore not be representative of broader sex effects on these connectivity measures, as there could be a selection bias or an uncontrolled effect of examining a sample of predominantly older individuals.

### Future Directions

Future work should also conduct more detailed analyses of specific regional connectivity-cognition associations in brain circuits which are structurally or functionally implicated in the cognitive processes of interest. For example, local connectivity of the left temporal lobe, the right posterior parietal lobe, and the dorsolateral prefrontal cortex can be compared by sex and individually related to differences in verbal memory, spatial memory, or executive functioning respectively (Äikiä et al. 2001; Berryhill and Olson 2008; Stuss 2011). Different global and local connectivity measures should be investigated and compared to see if the particular measure selected has implications related to the relevant cognitive task at hand (Imms et al. 2021). It may also be interesting to examine differences in intrahemispheric and interhemispheric connectivity, to see if, as found by Ingalhalikar and colleagues (Ingalhalikar et al. 2014), females will tend to show more interhemispheric connectivity and males more intrahemispheric connectivity, and if this is related to differences in cognitive performance.

Studies stemming from this work should also examine other morphological measures, such as surface area and volume. Examining volumes in particular would allow for the inclusion of many subcortical regions (e.g., hippocampus), which are often excluded in surface-based processing pipelines, such as CIVET. Examining these highly-implicated areas of the brain for both verbal memory and spatial memory might be informative for understanding the differences which underlie cognitive performance (Bonner-Jackson et al. 2015). Furthermore, multimodal brain connectivity analyses, such as analyses that combine structural neuroimaging data with DTI tractography (e.g. Gong et al. 2009) or functional MRI (fMRI) data (e.g. Liu et al. 2008; Meda et al. 2009) could also provide greater insight into how these different connectivity measures map onto each other, and thereby how graph-based structural connectivity findings should be better interpreted.

In addition to the general population, it would be interesting to examine how sex differences in cognition and brain connectivity translate to clinical populations. Differences in structural brain connectivity have previously been related to cognitive sex differences in clinical samples. For example, in a study using regional brain volumes, Abbs and colleagues (Abbs et al. 2011) found sex differences in the structural covariance of verbal memory circuitry for patients with schizophrenia, but not healthy controls. Interestingly, among patients, these differences in structural covariance-derived connectivity were significantly predictive of sex differences in verbal memory performance. Furthermore, Buck and colleagues (Buck et al. 2022) observed that the poorer verbal memory demonstrated by male first-episode psychosis patients was associated with sex-dependent circuitry alterations. Future studies, such as those using patient data from the UKBB, can further investigate the way these sex differences may be moderated in clinical populations.

## Conclusion

In this study, we reproduced the well-established male spatial advantage and female verbal advantage in cognition. Female sex was additionally associated with faster processing speed. Global efficiency, as well as nodal strengths in both hemispheres, were found to be higher in males than in females, suggesting that males may have both higher global and local brain connectivity. Higher overall connectivity in males as indicated by global efficiency was found to mediate all observed sex differences in cognition, predicting poorer verbal memory performance, better spatial memory, and slower processing speed in males. Greater investigation into how sex differences in cognition and brain connectivity affect the association between brain connectivity and cognitive performance is needed. Findings from the present study will help contribute to an improved understanding of the way biological sex and differences in cognitive performance are related to measures of structural brain connectivity as derived from structural graph-theoretic methods.

## Acknowledgement

This study used the NeuroHub infrastructure and was undertaken thanks in part to funding from the Canada First Research Excellence Fund, awarded through the Healthy Brains, Healthy Lives initiative at McGill University. This research was enabled in part by support provided by Calcul Québec and Compute Canada, and has been conducted using the UK Biobank Resource under Application Number 45551. Salary awards include a James McGill professorship to ML, a Mitacs Accelerate fellowship in partnership with Otsuka Canada to KML, and funding through Canadian Institutes of Health Research (CIHR #183720) to JFT.

## Statement of Contribution

The study was conceived and designed by CCY, KML, and ML. The identification of background papers was performed by CCY with guidance from KML. Data from the UKBB repository was collected, entered, and coded by KML, J. Unrau, D. Mendelson, and CCY. Inhouse code was developed by KML, JFT, and CCY. Results were analyzed and interpreted by CY with the input of KML and ML. The manuscript was prepared by CY, with draft revisions made according to the recommendations of KML, JFT, and ML. The final manuscript was reviewed and approved by ML.

