## Supplementary Figure 1 for "Normative Sex Differences in Cognition and Morphometric Brain Connectivity: Evidence from 30,000+ UK Biobank Participants"

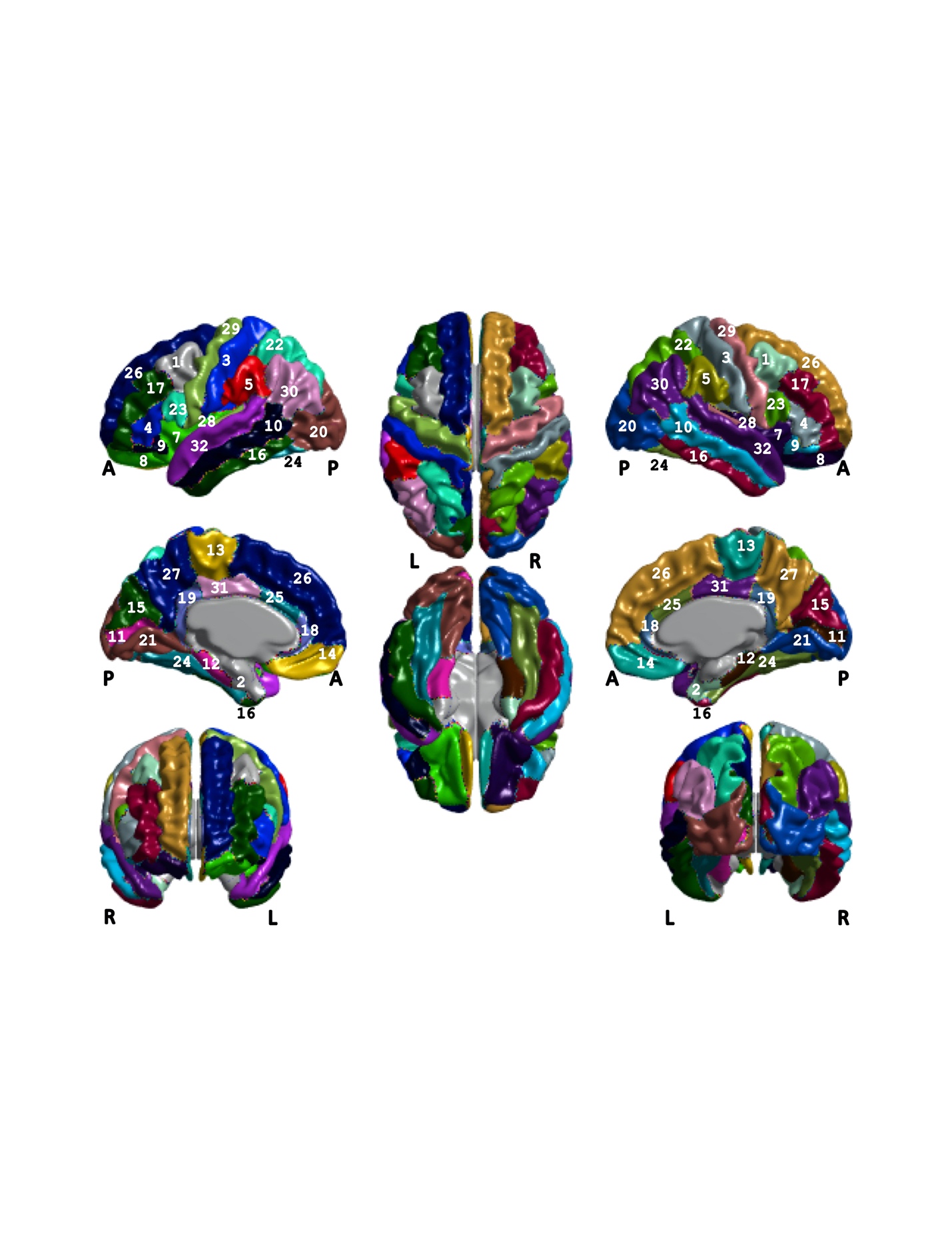

Figure S1. Desikan-Killiany-Tourville (DKT) parcellation. **A:** anterior, **L:** left, **P:** posterior, **R:** right, **1:** caudal middle frontal, **2:** entorhinal cortex, **3:** postcentral gyrus, **4:** lateral frontal triangularis, **5:** supramarginal gyrus, **7:** insula, **8:** lateral orbitofrontal, **9:** lateral frontal orbitalis, **10:** middle temporal, **11:** pericalcarine, **12:** parahippocampal, **13:** paracentral gyrus, **14:** medial orbitofrontal, **15:** cuneus, **16:** inferior temporal, **17:** rostral middle frontal, **18:** rostral anterior cingulate, **19:** isthmus cingulate gyrus, **20:** lateral occipital, **21:** lingual gyrus, **22:** superior parietal, **23:** lateral frontal opercularis, **24:** fusiform gyrus, **25:** caudal anterior cingulate, **26:** superior frontal gyrus, **27:** precuneus, **28:** transverse temporal, **29:** precentral gyrus, **30:** inferior parietal, **31:** posterior cingulate, **32:** superior temporal.
