## Supplementary Figure 1 for "Normative Sex Differences in Cognition and Morphometric Brain Connectivity: Evidence from 30,000+ UK Biobank Participants"

Supplementary Figure 2

*Hypothesized Mediation Model of Sex, Global Efficiency, and Cognition*

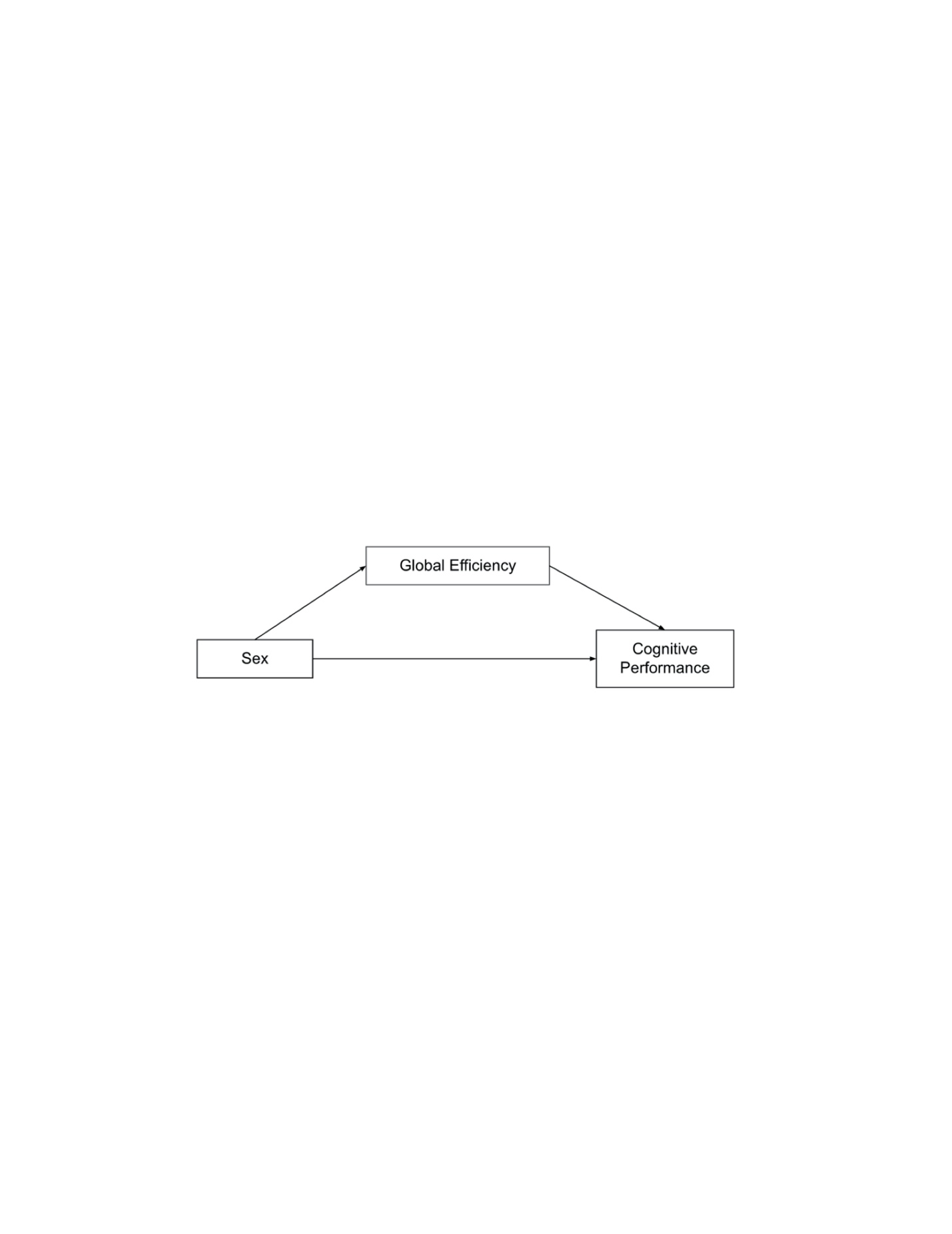

*Note.* This figure corresponds to Model 4 in PROCESS macro (Hayes, 2018).
