## Supplemental Table 1 for "Normative Sex Differences in Cognition and Morphometric Brain Connectivity: Evidence from 30,000+ UK Biobank Participants"

Supplementary Table 1. Desikan-Killiany-Tourville regions exhibiting higher morphometric brain connectivity strengths in males.

| Region | *F*(2,31077) | *p(F)* | $\boldsymbol{R}_{\boldsymbol{adj}}^{\boldsymbol{2}}$ | B | *T* | *p(t)* | *R^2^* |
| --- | --- | --- | --- | --- | --- | --- | --- |
| Precuneus L | 337.0 | < .001 | .02 | 6.66 | 18.36 | < .001 | .02 |
| Superior Parietal L | 321.7 | < .001 | .02 | 6.41 | 17.93 | < .001 | .02 |
| Paracentral Gyrus L | 319.4 | < .001 | .02 | 6.36 | 17.21 | < .001 | .02 |
| Superior Parietal R | 344.3 | < .001 | .02 | 6.19 | 17.19 | < .001 | .02 |
| Inferior Occipital Cortex R | 225.9 | < .001 | .01 | 5.95 | 16.39 | < .001 | .01 |
| Transverse Temporal L | 175.6 | < .001 | .01 | 5.67 | 15.67 | < .001 | .01 |
| Isthmus Cingulate Gyrus R | 222.0 | < .001 | .01 | 5.38 | 15.05 | < .001 | .01 |
| Precuneus R | 248.2 | < .001 | .02 | 5.35 | 14.61 | < .001 | .02 |
| Lingual Gyrus L | 157.7 | < .001 | .01 | 5.31 | 14.82 | < .001 | .01 |
| Inferior Occipital Cortex L | 190.2 | < .001 | .01 | 5.3 | 14.54 | < .001 | .01 |
| Supramarginal Gyrus L | 181.4 | < .001 | .01 | 5.23 | 14.36 | < .001 | .01 |
| Inferior Parietal L | 286.3 | < .001 | .02 | 5.22 | 14.46 | < .001 | .02 |
| Paracentral Gyrus R | 292.3 | < .001 | .02 | 5.16 | 13.93 | < .001 | .02 |
| Superior Frontal Gyrus R | 134.4 | < .001 | .01 | 5.09 | 14.55 | < .001 | .01 |
| Lateral Orbitofrontal L | 115.0 | < .001 | .01 | 5.00 | 12.49 | < .001 | .01 |
| Cuneus R | 154.4 | < .001 | .01 | 5.00 | 14.06 | < .001 | .01 |
| Cuneus L | 162.8 | < .001 | .01 | 4.99 | 14.13 | < .001 | .01 |
| Parahippocampal L | 189.1 | < .001 | .01 | 4.97 | 14.73 | < .001 | .01 |
| Isthmus Cingulate Gyrus L | 200.5 | < .001 | .01 | 4.92 | 14 | < .001 | .01 |
| Transverse Temporal R | 138.4 | < .001 | .01 | 4.81 | 13.17 | < .001 | .01 |
| Fusiform Gyrus R | 137.4 | < .001 | .01 | 4.80 | 13.65 | < .001 | .01 |
| Parahippocampal R | 157.6 | < .001 | .01 | 4.76 | 13.51 | < .001 | .01 |
| Lingual Gyrus R | 112.6 | < .001 | .01 | 4.62 | 13.2 | < .001 | .01 |
| Posterior Cingulate L | 191.9 | < .001 | .01 | 4.49 | 12.69 | < .001 | .01 |
| Fusiform Gyrus L | 138.9 | < .001 | .01 | 4.43 | 12.23 | < .001 | .01 |
| Pericalcarine L | 169.5 | < .001 | .01 | 4.39 | 12.08 | < .001 | .01 |
| Superior Temporal L | 145.0 | < .001 | .01 | 4.39 | 12.19 | < .001 | .01 |
| Postcentral Gyrus L | 182.7 | < .001 | .01 | 4.24 | 12.09 | < .001 | .01 |
| Posterior Cingulate R | 158.1 | < .001 | .01 | 4.14 | 11.71 | < .001 | .01 |
| Superior Frontal Gyrus L | 117.0 | < .001 | .01 | 4.02 | 11.53 | < .001 | .01 |
| Superior Temporal R | 88.0 | < .001 | .01 | 3.97 | 9.7 | < .001 | .01 |
| Caudal Middle Frontal L | 236.7 | < .001 | .02 | 3.93 | 10.61 | < .001 | .02 |
| Pericalcarine R | 108.8 | < .001 | .01 | 3.91 | 11.01 | < .001 | .01 |
| Middle Temporal R | 100.3 | < .001 | .01 | 3.84 | 8.94 | < .001 | .01 |
| Rostral Middle Frontal L | 279.5 | < .001 | .02 | 3.81 | 10.26 | < .001 | .02 |
| Postcentral Gyrus R | 156.5 | < .001 | .01 | 3.77 | 10.8 | < .001 | .01 |
| Lateral Frontal Orbitalis L | 63.13 | < .001 | .00 | 3.66 | 9.73 | < .001 | .00 |
| Lateral Orbitofrontal R | 105.5 | < .001 | .01 | 3.64 | 10.04 | < .001 | .01 |
| Supramarginal Gyrus R | 91.7 | < .001 | .01 | 3.57 | 8.97 | < .001 | .01 |
| Caudal Middle Frontal R | 197.0 | < .001 | .01 | 3.54 | 9.73 | < .001 | .01 |
| Inferior Parietal R | 127.7 | < .001 | .01 | 3.49 | 7.7 | < .001 | .01 |
| Rostral Middle Frontal R | 200.7 | < .001 | .01 | 3.40 | 9.32 | < .001 | .01 |
| Precentral Gyrus R | 206.2 | < .001 | .01 | 3.38 | 8.76 | < .001 | .01 |
| Lateral Frontal Triangularis L | 131.8 | < .001 | .01 | 3.37 | 9.37 | < .001 | .01 |
| Caudal Anterior Cingulate R | 189.0 | < .001 | .01 | 3.30 | 8.74 | < .001 | .01 |
| Lateral Frontal Triangularis R | 102.9 | < .001 | .01 | 3.24 | 8.76 | < .001 | .01 |
| Medial Orbitofrontal R | 95.7 | < .001 | .01 | 3.23 | 8.5 | < .001 | .01 |
| Rostral Anterior Cingulate R | 192.7 | < .001 | .01 | 3.04 | 8.16 | < .001 | .01 |
| Caudal Anterior Cingulate L | 156.4 | < .001 | .01 | 2.92 | 8.26 | < .001 | .01 |
| Lateral Frontal Opercularis R | 60.9 | < .001 | .00 | 2.91 | 7.87 | < .001 | .00 |
| Middle Temporal L | 120.8 | < .001 | .01 | 2.89 | 7.87 | < .001 | .01 |
| Lateral Frontal Opercularis L | 68.0 | < .001 | .00 | 2.86 | 7.89 | < .001 | .00 |
| Lateral Frontal Orbitalis R | 49.5 | < .001 | .00 | 2.75 | 7.46 | < .001 | .00 |
| Medial Orbitofrontal L | 55.8 | < .001 | .00 | 2.55 | 6.75 | < .001 | .00 |
| Rostral Anterior Cingulate L | 134.7 | < .001 | .01 | 2.26 | 6.07 | < .001 | .01 |
| Inferior Temporal R | 83.5 | < .001 | .01 | 2.25 | 6.13 | < .001 | .01 |
| Entorhinal Cortex R | 69.1 | < .001 | .00 | 2.14 | 5.89 | < .001 | .00 |
| Inferior Temporal L | 39.2 | < .001 | .00 | 1.56 | 4.48 | < .001 | .00 |
| Precentral Gyrus L | 236.6 | < .001 | .02 | 0.05 | 7.98 | < .001 | .02 |
| Entorhinal Cortex L | 62.7 | <.001 | .00 | 0.76 | 2.07 | .038 | .00 |
| Insula L | 47.15 | <.001 | .00 | 0.18 | 0.50 | .615 | .00 |
| Insula R | 7.5 | <.001 | .00 | 0.01 | 0.02 | .982 | .00 |

*Note.* Region = DKT region, SE = standard error, $R_{adj}^{2}$= adjusted R squared, *R^2^* = multiple R squared, L = Left, R = Right, *p(t)* = Bonferroni-corrected p value.
